# Radio-pathomic maps of cell density identify glioma invasion beyond traditional MR imaging defined margins

**DOI:** 10.1101/2021.04.07.438823

**Authors:** Samuel A. Bobholz, Allison K. Lowman, Michael Brehler, Fitzgerald Kyereme, Savannah R. Duenweg, John Sherman, Sean McGarry, Elizabeth J. Cochran, Jennifer Connelly, Wade M. Mueller, Mohit Agarwal, Anjishnu Banerjee, Peter S. LaViolette

## Abstract

Current MRI signatures of brain cancer often fail to identify regions of hypercellularity beyond the contrast enhancing region. Therefore, this study used autopsy tissue samples aligned to clinical MRIs in order to quantify the relationship between intensity values and cellularity, as well as to develop a radio-pathomic model to predict cellularity using MRI data. This study used 93 samples collected at autopsy from 44 brain cancer patients. Tissue samples were processed, stained for hematoxylin and eosin (HE) and digitized for nuclei segmentation and cell density calculation. Pre- and post-gadolinium contrast T1-weighted images (T1, T1C), T2 fluid-attenuated inversion recovery (FLAIR) images, and apparent diffusion coefficient (ADC) images calculated from diffusion imaging were collected from each patients’ final acquisition prior to death. In-house software was used to align tissue samples to the FLAIR image via manually defined control points. Mixed effect models were used to assess the relationship between single image intensity and cellularity for each image. An ensemble learner was trained to predict cellularity using 5 by 5 voxel tiles from each image, employing a 2/3-1/3 train-test split for validation. Single image analyses found subtle associations between image intensity and cellularity, with a less pronounced relationship within GBM patients. The radio-pathomic model was able to accurately predict cellularity in the test set (RMSE = 1015 cells/mm2) and identified regions of hypercellularity beyond the contrast enhancing region. We concluded that a radio-pathomic model for cellularity is able to identify regions of hypercellular tumor beyond traditional imaging signatures.

## 1. Introduction

Brain cancer, along with other central nervous system cancers, are the tenth leading cause of death worldwide, with an estimated 5-year survival rate of approximately 38 percent^1^. In particular, high-grade primary brain tumors such as glioblastomas are associated with particularly dismal prognoses, with a mean survival rate of around 12-18 months post-diagnosis^2^. Current standard of care for tumor patients includes targeted surgical resection of the tumor area, as defined by magnetic resonance imaging (MRI), targeted radiation therapy, and administration of chemotherapeutic agents. At recurrence, patients can be treated with salvage therapy such as bevacizumab^3,4^, with many patients also opting for newer FDA-approved treatments such as tumor-treating fields^5–7^. Precise localization of tumor margins is essential to maximizing the efficacy of these treatments and monitoring tumor progression.

Magnetic resonance imaging (MRI) is currently the gold standard for identifying the tumor boundary and monitoring disease progression. T1-weighted imaging acquired following an injection with a gadolinium contrast agent (T1C) is used to identify regions where active tumor has disrupted the blood-brain barrier. Contrast enhancement is often used to define the extent of the primary tumor region^8^. Hyperintense regions on fluid-attenuated inversion recovery (FLAIR) images are thought to indicate a combination of tumor-related edema^9–11^ and infiltrative non-enhancing tumor^12^. Multi-b-value diffusion weighted imaging (DWI) is also typically included in glioma imaging protocols, which is used to calculate quantitative apparent diffusion coefficient (ADC) maps. These maps identify areas of restricted diffusion that may indicate either hypercellular tumor^13–16^ or coagulative necrosis^17^.

Critical for understanding the relationship between radiological signatures and pathological features of the tumor is tissue-based pathological validation. These studies inherently require invasive tissue sampling, and therefore have typically been limited to surgical biopsy cores taken in-vivo from contrast-enhancing regions (i.e. suspected tumor). Studies using biopsy cores have provided pathological validation for imaging signatures such as the inverse relationship between ADC and cellularity^13,18,19)^. Combined radiological-pathological (rad-path) datasets have also been used to develop predictive models for pathological features^20–23^. While these studies have provided understanding of the relationship between the tumor environment and its corresponding radiological features, limitations in the number and size of tissue samples result in a loss of the ability to fully characterize these heterogeneous tumors.

Tumor heterogeneity has been a recent focus in radiology imaging studies of GBM. With non-invasive imaging, regional heterogeneity is readily measurable, but capturing heterogeneity with pathology samples is challenging. It can be achieved with en-bloc resection^24,25^ or sampling multiple times with image guided biopsies. Each of these techniques require properly aligning samples to their location, which can be difficult due to a lack of orientation. Other issues such as brain shift during craniotomy, and the inability to sample regions outside the suspected tumor region further complicate these strategies. Despite these challenges, pathological measurement of tumor heterogeneity is crucial to improving the localization of multiple tumor pathologies, as well as in order to validate imaging signatures beyond the currently accepted tumor boundary. In particular, high-grade gliomas are typically associated with diverse histopathological features that may confound traditional imaging interpretations, therefore validation across a range of suspected tumor characteristics are important to distinguish diagnostic differences in MR interpretation.

Another factor in the field of glioma rad-path correlation is the timing of tissue acquisition. Because the majority of surgeries and biopsies are done at the beginning of treatment, prior to radiation and chemotherapy, the imaging characteristics at that stage may not fully capture the spectrum of tissue, which evolves during treatment. Due to factors such as post-surgical blood products, radiation necrosis, and pseudo-progression, pathological correlations found upfront may not generalize in the treated phase^26–28^.

Studies of glioma patients at autopsy have shown that viable tumor can exist as far as 10 cm beyond contrast enhancement, where pseudo-palisading necrosis and other heterogenous pathological features may confound traditional MRI interpretations^15,17,29^. Due to the sampling limitations of biopsy samples, further pathological validation of MR imaging signatures is warranted, both beyond the contrast enhancing region and in the post-treatment state. Therefore, this study used large format tissue samples collected across the whole brain at autopsy in order to validate current imaging signatures, as well as use this pathological ground truth to develop predictive tools to assess prospective tumor beyond the contrast enhancing region. Specifically, this study tested the hypotheses that 1) MP-MRI intensity values are associated with tumor cellularity at autopsy 2) cellularity imaging signatures are more robust in non-GBM patients than GBM patients and 3) a radio-pathomic model trained on autopsy data will be able to accurately identify regions of hypercellular tumor beyond traditional imaging signatures.

## 2. Methods

### 2.1. Patient Population

A total of 44 patients with pathologically confirmed brain tumors were enrolled in this IRB-approved study. Patients underwent an autopsy following clinical decline and death. Clinical histories and demographic information are shown in **Table 1**. A diagrammatic representation of the tissue and imaging data collection process is shown in **Figure 1**.

**Figure 1:**
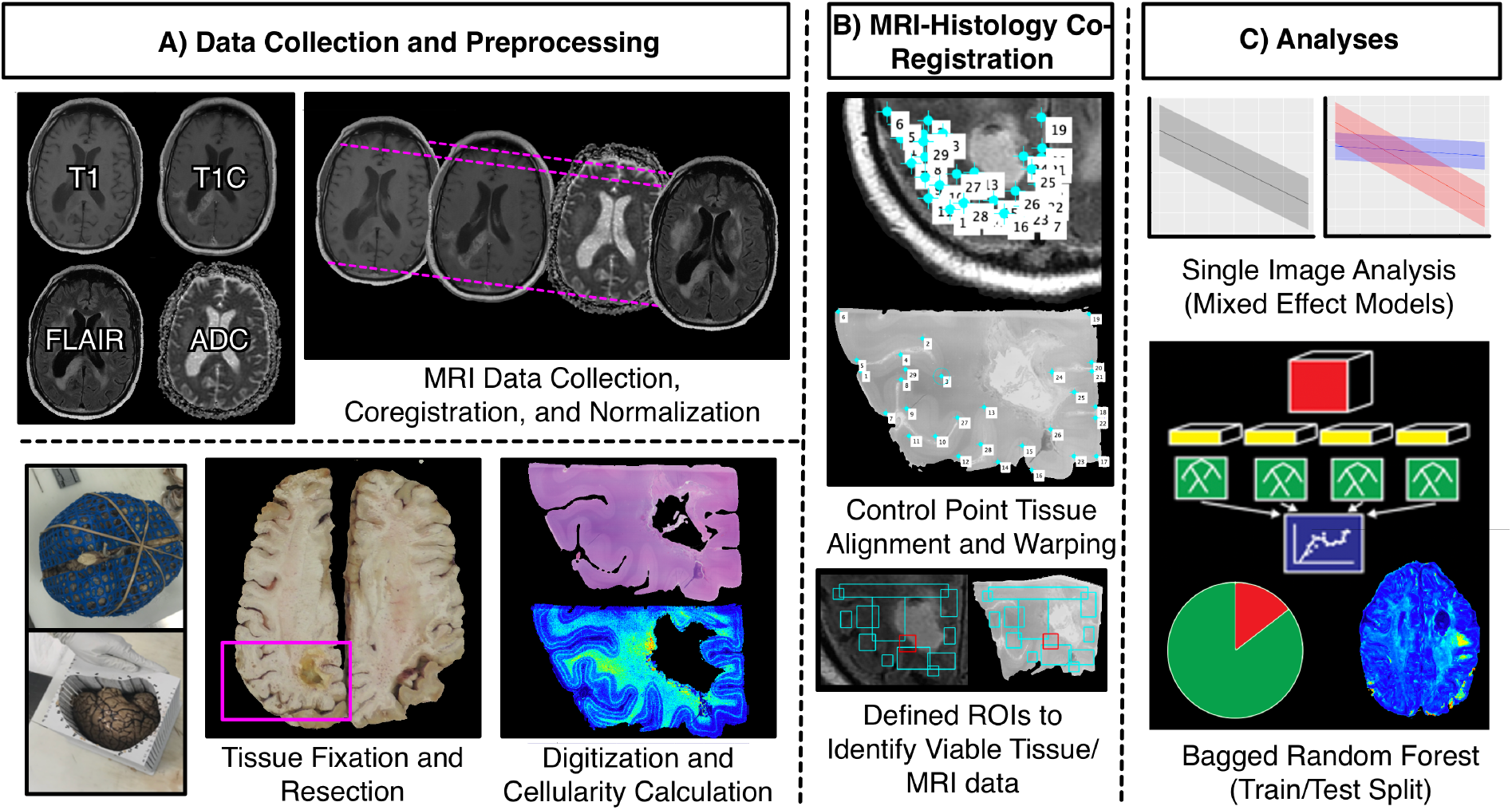
Overview of the data collection process. A) MRI data is collected from the patients’ final imaging session prior to death, co-registered, and T1, T1C, and FLAIR images are intensity normalized. Tissue fixation and sampling involves the use of 3D printed brain cages and slicing jigs in order to preserve structural integrity relative to the MRI. Following staining, tissue samples are digitized for cellularity calculation using an automated nuclei segmentation algorithm. B) In-house software is used to align each tissue sample to the FLAIR image using manually-defined control points and regions of interest. C) Single-image cellularity associations were computed using mixed effect models, and a bagging regression ensemble was trained to predict cellularity using 5 by 5 voxel tiles from each image using a 2/3-1/3 train-test split.

**Table 1:**
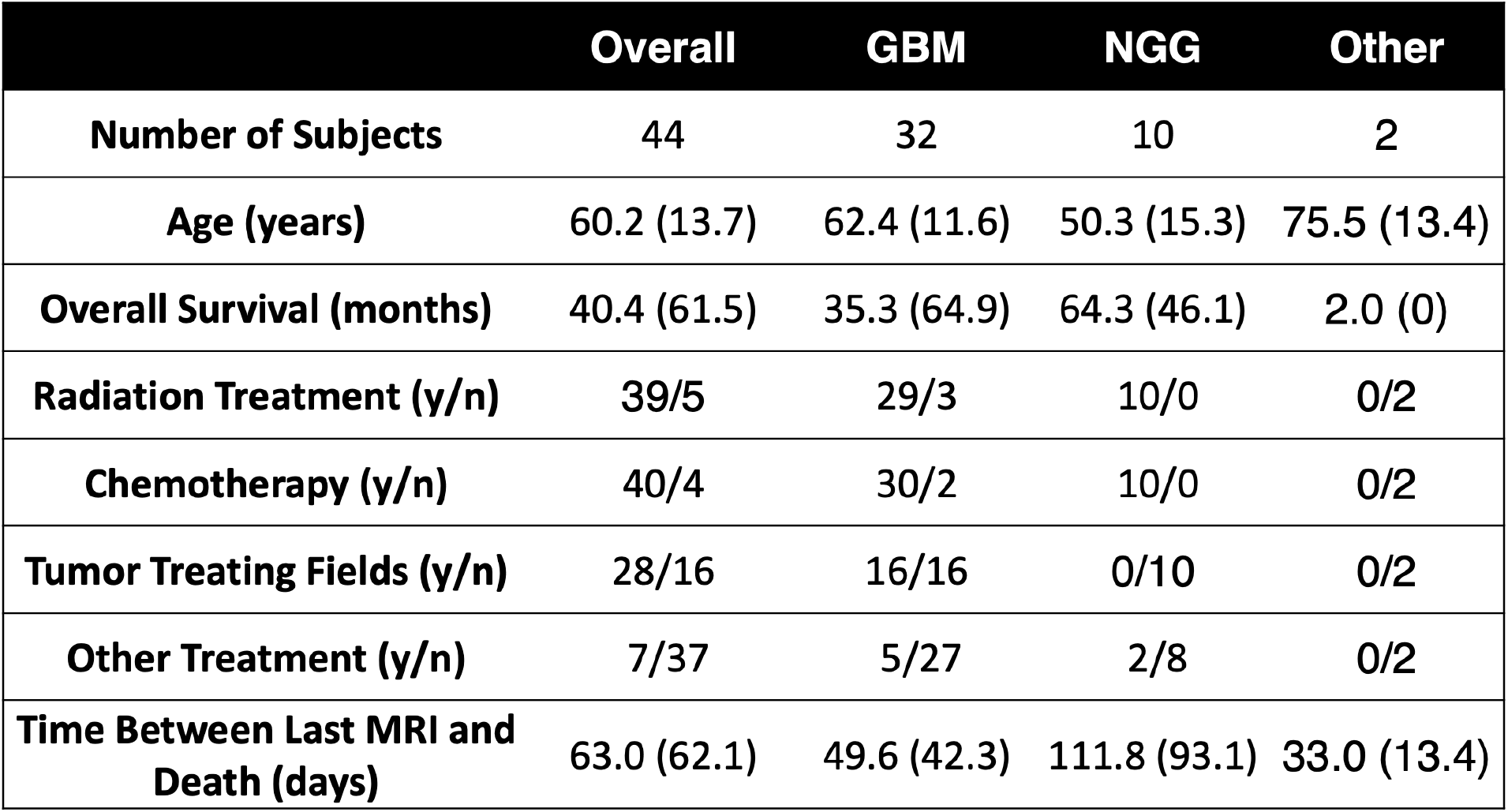
Clinical and demographic summary for study sample. Quantitative values are presented as mean (standard deviation).

|  | Overall | GBM | NGG | Other |
| --- | --- | --- | --- | --- |
| **Number of Subjects** | 44 | 32 | 10 | 2 |
| **Age (years)** | 60.2 (13.7) | 62.4 (11.6) | 50.3 (15.3) | 75.5 (13.4) |
| **Overall Survival (months)** | 40.4 (61.5) | 35.3 (64.9) | 64.3 (46.1) | 2.0 (0) |
| **Radiation Treatment (y/n)** | 39/5 | 29/3 | 10/0 | 0/2 |
| **Chemotherapy (y/n)** | 40/4 | 30/2 | 10/0 | 0/2 |
| **Tumor Treating Fields (y/n)** | 28/16 | 16/16 | 0/10 | 0/2 |
| **Other Treatment (y/n)** | 7/37 | 5/27 | 2/8 | 0/2 |
| **Time Between Last MRI and Death (days)** | 63.0 (62.1) | 49.6 (42.3) | 111.8 (93.1) | 33.0 (13.4) |

### 2.2. MR Image Acquisition and Preprocessing

The clinical MRI scans acquired closest to each patient’s death were used for this study. Each protocol included T1-weighted pre- and post-contrast images (T1 and T1C, respectively), T2-weighted fluid-attenuated inversion recovery images (FLAIR), and diffusion weighted images (DWI). Image acquisition was performed on our institutional MRI scanners, including 1.5T and 3T GE (General Electric Health, Waukesha, Wisconsin) and Siemens magnets (Siemens Healthineers, Erlangen, Germany). Example acquisition parameters for scans collected at 1.5T include (repetition time/echo time): T1 spin-echo sequence (T1), 666/14 ms; contrast-enhanced T1 acquired with gadolinium (T1C), 666/14 ms; apparent diffusion coefficient (ADC), calculated from diffusion-weighted images (DWI) acquired with a spin-echo echo-planar sequence, 10,000/90.7 ms; and FLAIR, acquired with an inversion recovery sequence, 10,000/151.8 ms and TI of 2,200 ms. Example acquisition parameters for scans collected at 3T include: T1 spin-echo sequence, 716.7/10 ms, T1C 716.7/10 ms, ADC from DWI acquired with a spin-echo echo-planar sequence, 8,000/83.1 ms, and FLAIR acquired with an inversion recovery sequence, 10,000/121.1. All images were acquired with submillimeter in-plane resolution.

The images collected for inclusion in validation analyses included the T1, T1C, and FLAIR images, as well as the ADC images derived from the DWI. All images were rigidly aligned with each subject’s FLAIR image using FSL’s FLIRT tool^30–32^. All non-quantitative images (T1, T1C, FLAIR), were intensity normalized by dividing voxel intensity by its whole brain standard deviation^33,34^. Because ADC is a quantitative measure, these images were not intensity normalized.

### 2.3. Ex-vivo Histology Processing

At autopsy, each patient’s brain was removed and placed in a patient-specific 3D-printed brain mold, modeled from the patient’s most recent MRI. These molds were meant to prevent tissue distortion during 2 to 3 weeks of fixation in formalin (15,17,35). Following fixation, brains were sliced using patient-specific slicing jigs, also 3D printed to match axial slice orientation from the most recent MRI^17^. Tissue samples measuring approximately 2 inches by 3 inches were then collected from each subject from regions of suspected tumor, as well as tissue adjacent to the suspected tumor region. These samples were then processed, embedded in paraffin, sliced, and stained with hematoxylin and eosin (HE). The slides were then digitized at 0.2 microns per pixel (40X magnification) using a Huron sliding stage microscope. A total of 93 tissue samples were collected across all patients.

### 2.4. Pathological Feature Extraction

After digitization, images were processed using Matlab 2020b (Mathworks Inc., Natick, MA) in order to extract pathological features for quantitative analyses. First, a color deconvolution algorithm was used to project color data in terms of relative stain intensities, resulting in an image with color channels representing hematoxylin, eosin, and residual color information^36,37^. Images were then down sampled by a factor of 10 in order to smooth color data for improved nuclei segmentation, as well as to decrease processing time. Cell nuclei were highlighted by applying filters on each color channel to selectively identify positive hematoxylin staining, and individual nuclei were identified using Matlab’s regionprops function. Cell count was computed across 50 by 50 superpixels across the image and converted to cells per square millimeter for ease of interpretability. Additionally, segmentations for extra-cellular fluid (ECF) and cytoplasm were computed and converted to proportions of the superpixel occupied by the component of interest. All segmentations were visually inspected for quality assurance, and example segmentations are provided in **Supplemental Figure 1**.

### 2.5. MRI-Histology Co-registration

Previously published in-house software (Hist2MRI Toolbox, written in Matlab, Mathworks Inc, Natick, MA) was used to precisely align histology images to each patient’s clinical imaging^15,17,35,38,39^. Photographs taken during tissue sectioning were used to identify the imaging slice that corresponded to each tissue sample. Next, control points were manually defined in order to calculate a nonlinear warp to match each tissue sample to its shared anatomical features with the FLAIR image. Following warping, manually defined regions of interest are used to identify regions of both valid histological information (e.g. free from processing artifacts) and valid MRI information (e.g. free from motion artifacts). Voxel intensity values from T1, T1C, FLAIR, and ADC images, as well as cellularity values for each voxel were collected across these ROIs and used for subsequent analyses. Across all 93 samples, a total of 578,668 voxels were included

### 2.6. Statistical Analyses

#### 2.6.1. Single Image Analyses

Linear mixed effect models were used in order to quantify the propensity for individual images to identify regions of hypercellularity. Image intensity was included as a main effect, with time between last MRI scan and patient death (in days) and grey/white matter probability included as covariates. Patient number was included in the model to account for patient-specific confounds. Regression coefficients (values) and R-squared values were reported to quantify the relationship between MRI intensity and cellularity in terms of slope and explained variance, respectively. R-squared values were computed in terms of conditional and marginal effects as well, in order to specifically delineate the explained variance resulting from imaging signatures from that of subject-level variance.

In order to specifically compare diagnosis-level differences in the relationship between MRI intensity and cellularity, similar mixed effect models were computed for each image type including a term for the interaction between image intensity and diagnosis, with diagnostic groups corresponding to GBM, non-GBM glioma (NGG), and Other. The other category consisted of one patient with brain metastases and one patient with lymphoma. Both of these patients did not receive treatment for their brain tumors and can thus provide a proxy for rad-path relationships in the untreated state. Due to the large number of observations relative to the patient-level data set size, p-values were considered a poor measure of meaningful significance (all p¡0.00001). Therefore, measures of effect size are reported for this subanalysis. Analogous models were also calculated for extracellular fluid (ECF) and cytoplasm as the dependent variables in order to examine other cellular factors that may driving imaging values, which are presented in **Supplemental Figure 2**.

#### 2.6.2. Radio-Pathomic Modeling

A random forest ensemble algorithm was used as the framework for developing a radio-pathomic model of cellularity. Specifically, a bootstrap aggregating (bagging) model was used, which fits independent weak learners across several independent bootstrapped samples from the training data set in order to obtain a combined ensemble model that minimizes variance across learners^40,41^. Inputs for this model were intensity values from 5 by 5 voxel tiles across each image, in order to incorporate local spatial and contextual information. Model performance was assessed within the training dataset, across a leave-one-subject-out cross validation scheme. To test generalizability, the models were then applied to imaging data from an independent dataset from 14 subjects held-out of the training. Performance was quantified using root mean squared error (RMSE) values, which were standardized to the standard deviation of cellularity. Example predictions for each validation scheme were computed across the whole brain of each patient in order to assess the model’s ability to discern regions of hypercellularity beyond traditional imaging signatures. Additionally, cellularity prediction maps were computed for one subject across several clinical timepoints to evaluate the model’s ability to track tumor progression over the course of the disease state.

## 3. Results

### 3.1. Single-Image Analyses

Mixed effect model results for the single image analyses are presented in **Figure 2**. T1, T1C, and FLAIR images demonstrated positive associations with cellularity (T1 = 160.23 (5.11), Conditional R2 = 0.35, Marginal R2 = 0.012; T1C = 480.60(4.00), Conditional R2 = 0.44, Marginal R2 = −0.074; FLAIR = 152.50(3.12), Conditional R2 = 0.34, Marginal R2 = 0.010). ADC values demonstrated a negative association with cellularity (B = −153.72(5.45), Conditional R2 = 0.34, Marginal R2 = 0.008), however the strength of this relationship was much lower than expected. When splitting data by diagnostic group (GBM vs. NGG vs. Other), stronger relationships between image intensity and cellularity were observed for NGG and Other patients across each image except the pre-contrast T1, with the largest diagnostic discrepancy seen in the ADC-cellularity relationship (T1 = 135.56 (4.10), Conditional R2 = 0.15, Marginal R2 = 0.014; T1C = 615.22 (3.76), Conditional R2 = 0.30, Marginal R2 = 0.126; FLAIR = 277.51 (2.83), Conditional R2 = 0.18, Marginal R2 = 0.036; ADC = 168.24 (2.84), Conditional R2 = 0.16, Marginal R2 = 0.016). The GBM group showed an opposite direction relationship compared to the Other group for the pre-contrast T1 intensity, with only a subtle relationship seen in the NGG group.

**Figure 2:**
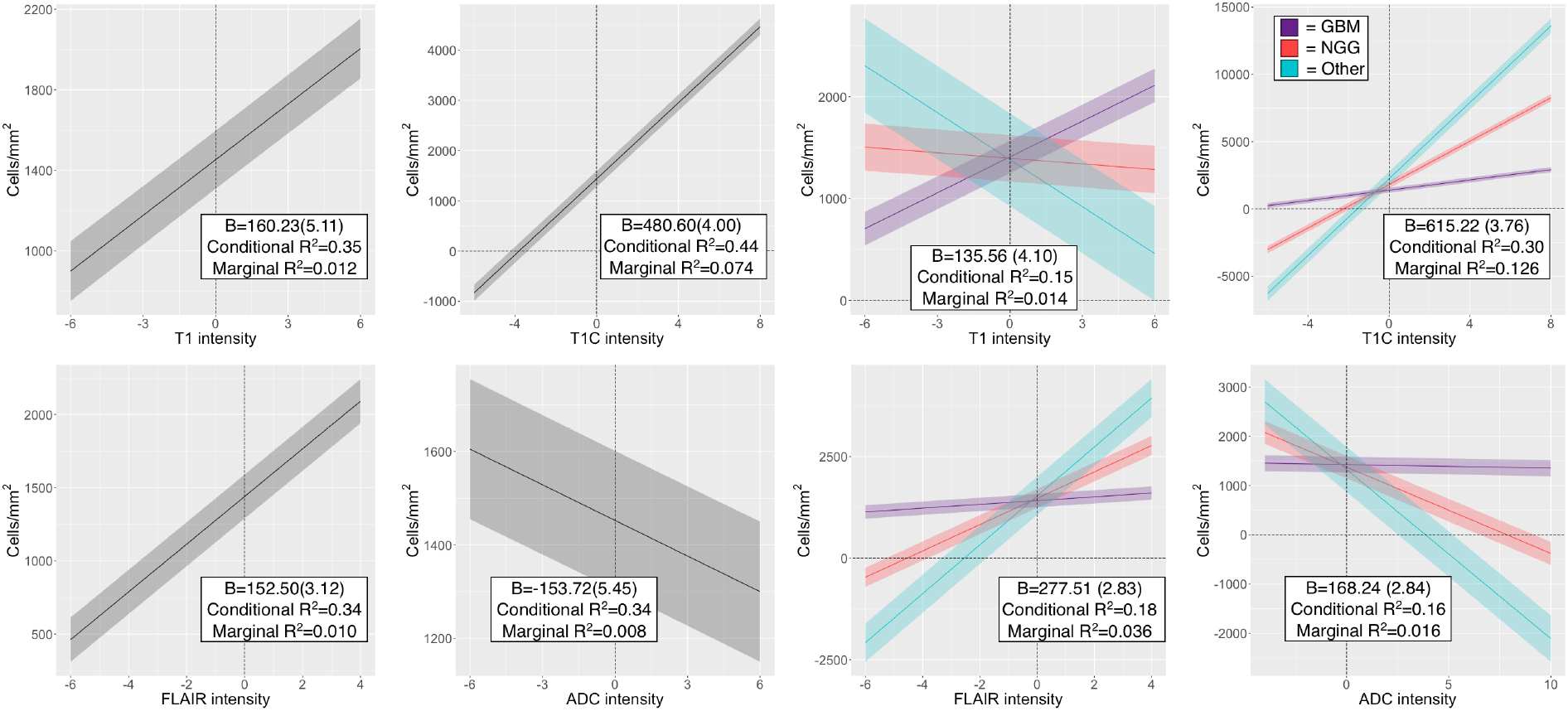
Single image results depicting the relationship between image intensity and cellularity for each contrast. values for the left-hand plots indicate the change in cellularity per standard deviation increase in image intensity, and indicate positive associations for T1, T1C, and FLAIR, with the expected negative association between ADC and cellularity present. B values for the right-hand plots indicate the difference in slope between GBM, NGG, and Other patients, indicating that GBM patients show less pronounced cellularity associations than non-GBM patients across all image types, with the exception of T1 intensity.

### 3.2. Radio-Pathomic Mapping

A summary of model performance, including train and test set root-mean-squared error values and a scatter plot summarizing example prediction values is provided in **Figure 3**. Overall train and test set RMSE values (389 cells/mm2 and 1015 cells/mm2, respectively) show some degree of overfitting with regards to the training data set, but generally indicate accurate prediction of cellularity. The scatterplot, which demonstrates example predictions in terms of T1SUB (T1C – T1), FLAIR, and ADC intensity values shows indications of expected relationships (i.e. FLAIR-ADC mismatch associated with hypercellularity), but also highlights that traditional hypercellularity signatures are often non-specific. Example predictions for test set subjects are included in **Figure 4**, where cellularity predictive maps (CPM) for the whole brain are provided with the clinical images for each patient, as well as the pathological ground truth from the aligned autopsy slide. These example predictions highlight that the CPMs calculated from the radio-pathomic model accurately predict several regions of increased cellularity beyond the traditional contrast enhancing region and discriminate between hypercellular and non-hypercellular regions within contrast enhancement. Additionally, example CPMs across longitudinal imaging for a 59-year-old male patient diagnosed with GBM are presented in **Figure 5** with timepoints just after initial surgery (511 days prior to death), as well as two imaging time points acquired during cognitive decline (26 and 12 days prior to death). These CPMs highlight the potential for the radio-pathomic model to monitor disease progression, as predictions begin to suggest selective proliferation within the contrast-enhancing region during the later stages of the disease.

**Figure 3:**
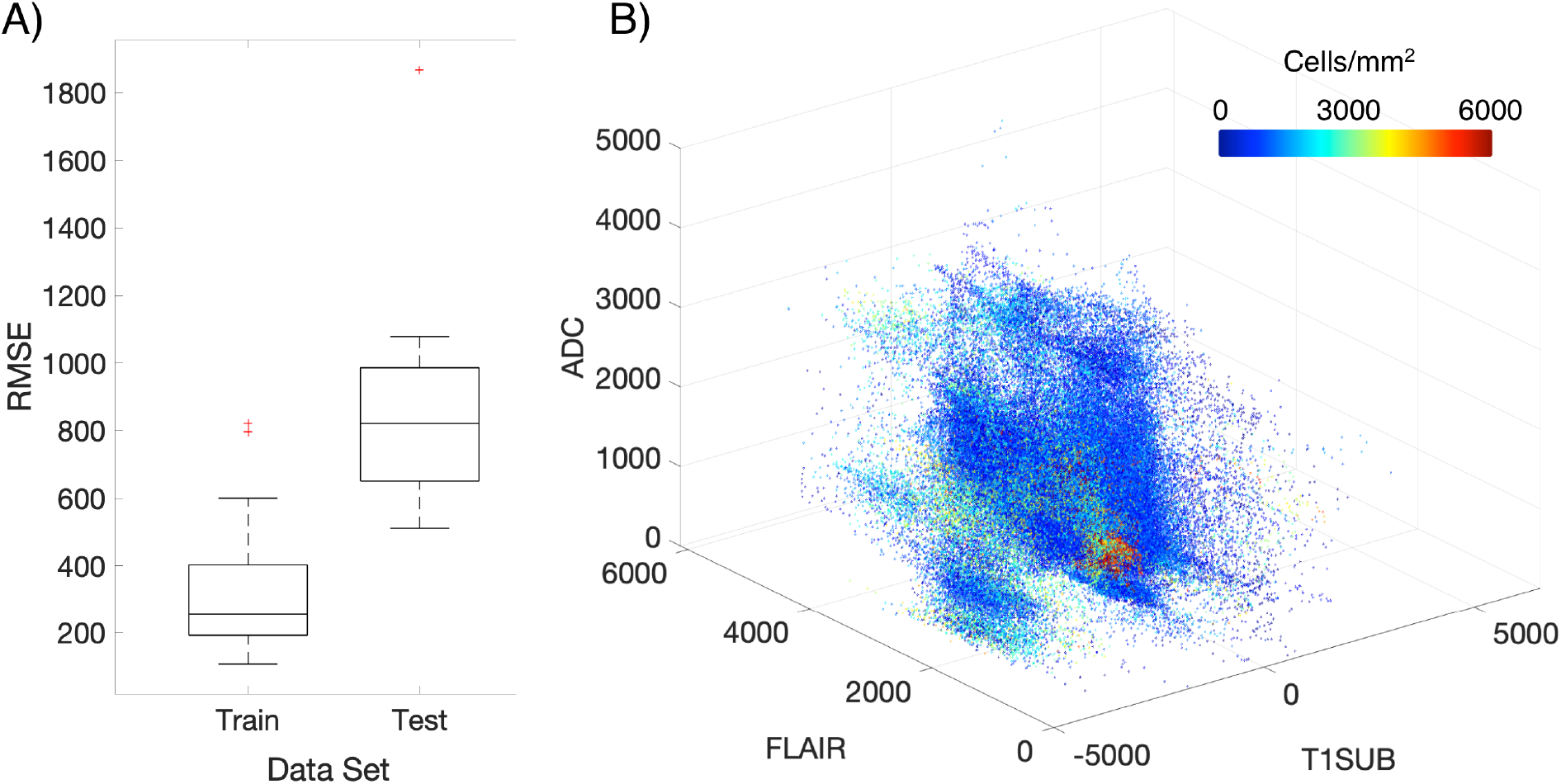
A) Subject-level RMSE values for the train and test data sets. Despite some degree of overfitting, the test set RMSE indicates that the radio-pathomic model is able to accurately predict cellularity across most subjects. B) Example predictions for test set imaging values presented in terms of their T1SUB, FLAIR, and ADC intensity values. Patterns suggest the presence of traditional imaging signatures, but also indicate the lack of specificity for these signatures with regards to hypercellularity

**Figure 4:**
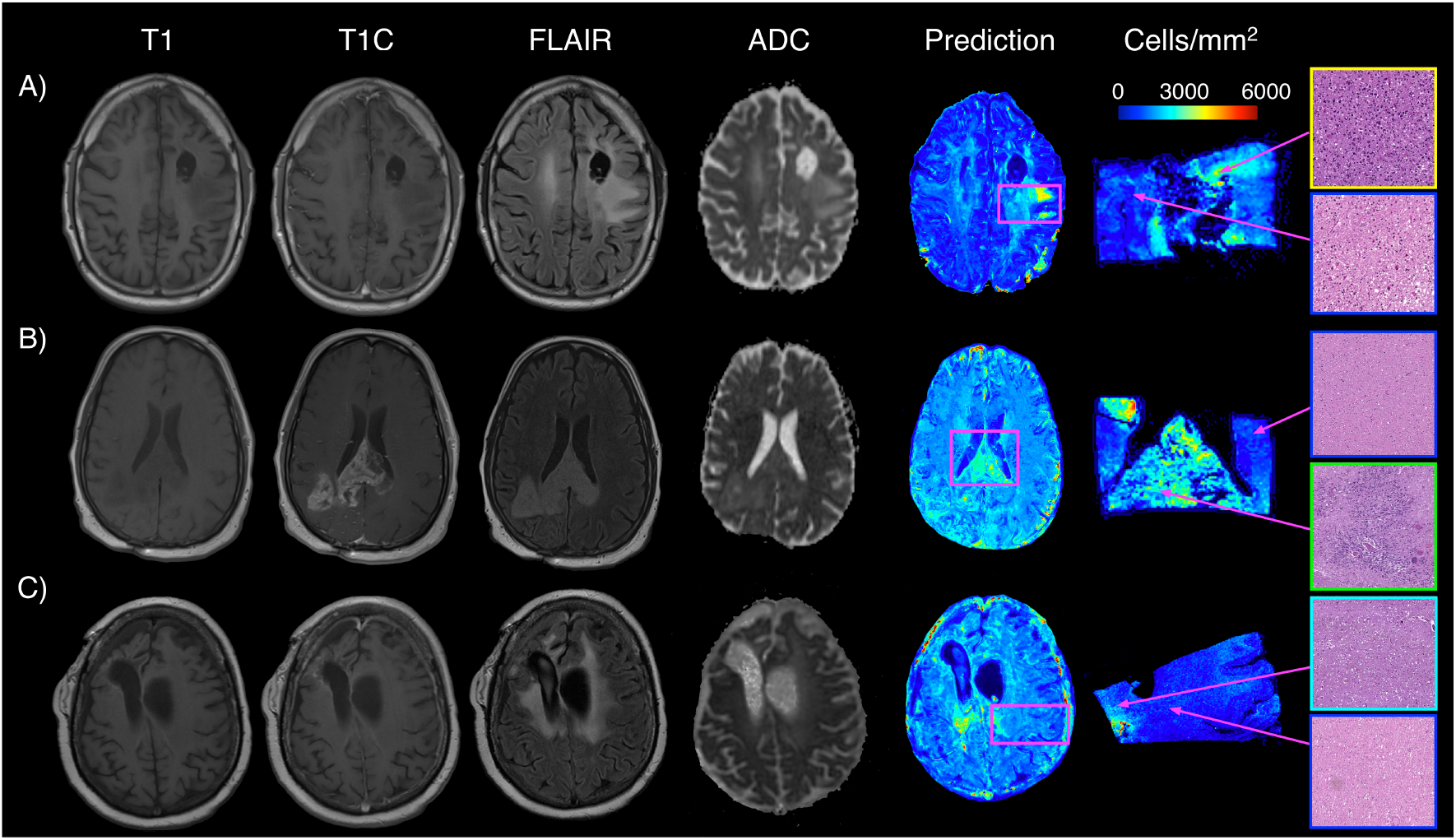
Example predictions for three representative subjects, including A) a 43-year-old male diagnosed with a grade 3 anaplastic astrocytoma, B) a 48 year old male diagnosed with a GBM, and C) a 31-year-old female diagnosed with a grade 3 anaplastic astrocytoma. These predictions indicate that the radio-pathomic model is able to predict regions of hypercellularity beyond the contrast-enhancing region, as well as in the absence of restricted diffusion on the ADC image

**Figure 5:**
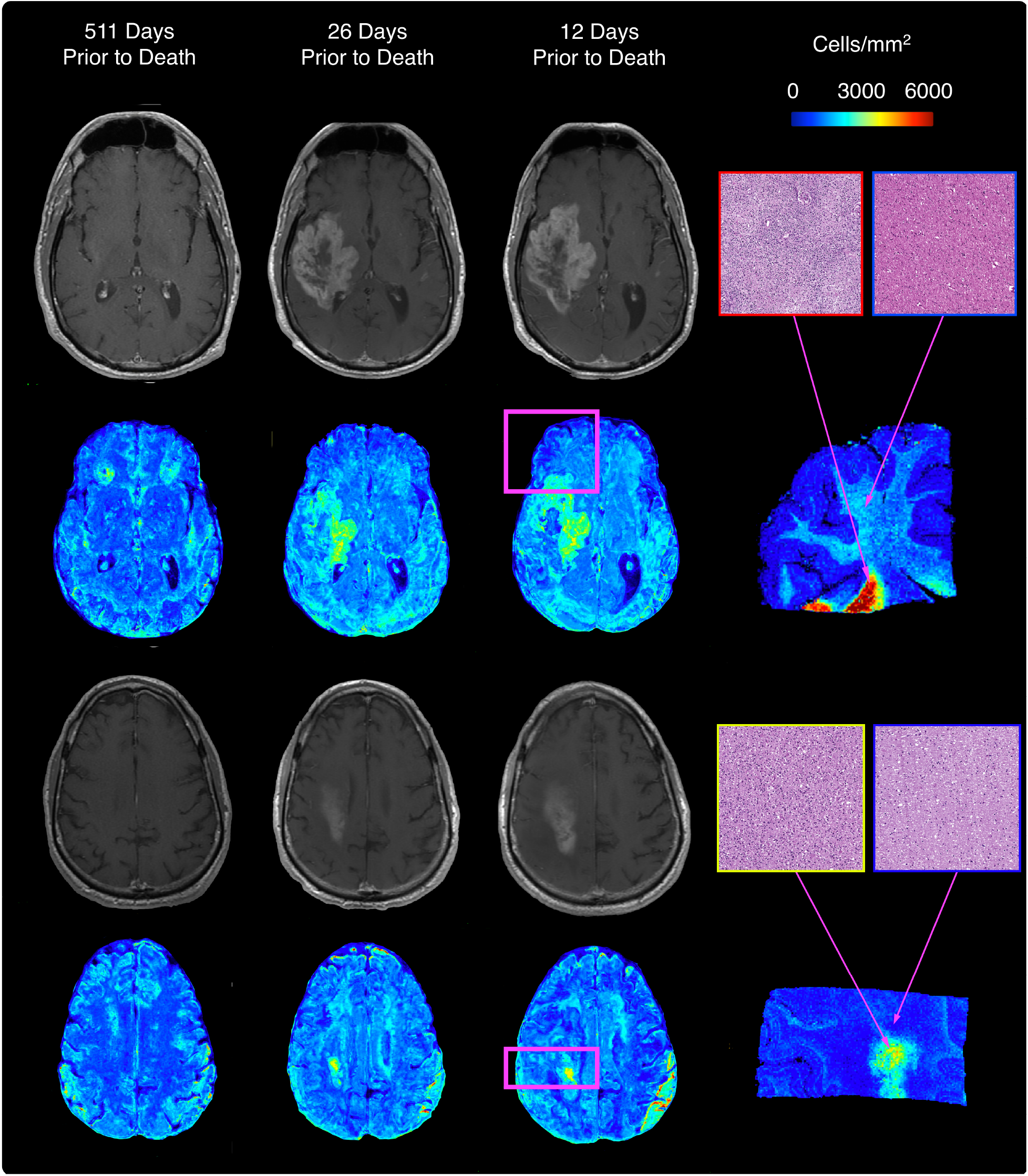
Longitudinal predictions for a 59-year-old male diagnosed with GBM at surgery. Earlier predictions indicate limited hypercellular tumor; however, the last two time points, collected around the beginning of cognitive decline, indicate hypercellular tumor growth within a portion of the contrast enhancing region.

## 4. Discussion

This study evaluated current MP-MRI signatures of brain tumor pathology in comparison to a radio-pathomic model for brain cancer. Using tissue samples taken at autopsy aligned to the patients’ clinical imaging, this study assessed relationships between imaging and pathology in the post-treatment state, as well as beyond the contrast-enhancing region. LME-based analyses of individual image intensity values found that single image signatures explain a relatively small proportion of cellularity variance. Additional single image analyses found an effect of tumor type on the cellularity-intensity relationship, with reduced cellularity associations seen in GBM compared to non-GBM patients across all image types. We developed a radio-pathomic model using a bagging ensemble architecture, which predicted cellularity accurately on withheld subjects, despite performing less reliably in a small subset of cases. Predictive maps showed that the model accurately predicted regions of hypercellularity beyond regions of contrast enhancement and other traditional imaging signatures, and predictions across longitudinal data plausibly tracked tumor proliferation over the course of the disease state.

Contrast-enhancement is often the principal MRI signature used to clinically define the tumor boundary^8,42^. Coupled with FLAIR hyperintense regions, these signatures are thought to indicate regions with some combination of infiltrative tumor and vasogenic edema^9,11^. The general trends for increased cellularity associated with increased contrast enhancement and FLAIR intensity support the notions that these features relate to pathological effects of the tumor. However, these features failed to account for the vast majority of cellularity variance, and in some cases failed to identify regions of hypercellular tumor beyond the primary tumor mass. Previous studies have suggested that radiation therapy and other treatments may influence the relationship between different imaging signatures and pathological tumor features, as induced necrosis may confound traditional interpretations of these features^43,44^. Diagnostic factors may play a role here as well, as the results of this study show that GBM cases, which present with a wide range of pathological characteristics, have less pronounced relationships between cellularity and imaging values than their lower-grade, more pathologically homogenous counterparts^45–47^. Further studies probing potential confounds could further assess which factors influence the relationship between T1C/FLAIR enhancement and its underlying pathological characteristics.

Past studies have particularly highlighted ADC values as a correlate for cellularity. Particularly, studies using biopsy tissue samples to calculate ground truth cellularity demonstrate a fairly robust inverse relationship between ADC and cellularity^13,16,48^. This study finds evidence of this negative association, though the strength of this relationship is more subtle in comparison to previous studies. Biopsy tissues sampled within regions of contrast enhancement may only reflect relationships near the tumor core, which may not generalize to areas beyond the traditionally defined tumor region. Additionally, diagnostic factors could play a role here, as a much more pronounced relationship between ADC and cellularity was observed for non-GBM cases compared to GBM cases. Heterogenous pathological features of GBM, such as pseudopalisading necrosis, may co-localize within individual voxels, mitigating the effects of cell density on diffusion restriction. Other pathological features may be more representative of ADC values as well, with both ECF and cytoplasm showing more consistent relationships across diagnostic groups (see Supplemental Figure 2). Given past findings examining the relationship between ADC and cellularity, further replications of these results are warranted, as well as more in-depth assessments of how heterogenous pathological findings relate to corresponding imaging signatures.

The performance statistics for our radio-pathomic model suggest that our model can accurately assess tumor cellularity in brain cancer patients. Despite some increased error observed in some individual cases, the majority of subjects in the test set had a RMSE within a standard deviation of each subjects’ cellularity, indicating that the model has the capacity to generalize to unseen data. These results thus demonstrate the feasibility of developing radio-pathomic models for pathological features using autopsy tissue data. These predictions are particularly valuable as they are validated in the post-treatment state, whereas predictive models from biopsy tissue are typically collected prior to treatments such as radiotherapy. Biopsy core studies also provide a much smaller region of tissue per subject, providing a single ground truth observation per sample. Autopsy tissue samples contain pathological substrates for hundreds of voxels per sample, providing a richer training data set for developing machine learning models.

In addition to traditional measures of model performance, we tested the hypothesis that a radio-pathomic model can highlight regions of tumor beyond contrast enhancement. Example whole-brain CPMs indicate that the radio-pathomic model can highlight regions of infiltrative tumor beyond contrast enhancement, suggesting that radio-pathomic modeling may provide improved localization of tumor areas in the post-treatment setting. In addition, CPMs for the longitudinal imaging demonstrate the potential for CPMs to continuously monitor disease progression and track the timing of non-contrast enhancing tumor infiltration. The CPMs generated from this radio-pathomic model provide early insight into clinical uses for non-invasive imaging models for pathological information collected at autopsy.

### 4.1. Limitations

While the results of this study are promising, there are several limitations that warrant noting. While this study uses a relatively large data set compared to other studies of brain cancer tissue at autopsy and developed voxel-wise predictions across hundreds of thousands of observations, the subject-level sample size is still fairly small by machine learning standards. Particularly for a disease with a wide range of pathological characteristics and prognoses, future investigations with a larger number of cases will be able to further improve the subject-level generalizability of radio-pathomic models. Additionally, the use of clinical imaging data provides another potential confound of these results, as variability in scanner vendor or image acquisition parameters may affect radio-pathomic relationships. Lastly, time between image acquisition and tissue sampling is an inherent concern arising from the use of autopsy tissue, which may affect these relationships beyond what we were able to control for statistically in this study. Future studies acquiring post-mortem MRI data may be able to assess the influence of this time period on assessing radio-pathomic relationships.

### 4.2. Conclusion

In conclusion, this study evaluated MP-MRI signatures of brain tumor pathology and developed a radio-pathomic model for brain cancer using machine learning. Our model defined predictive maps of tumor cellularity highlighted tumor beyond conventional boundaries and plausibly tracked tumor growth using longitudinal imaging. We hope these algorithms may be useful in the future for treatment planning and tumor monitoring. Additional research is necessary to probe the potential confounding factors influencing these predictions (diagnosis, treatment effects, etc.), as well as to further improve and validate these predictive maps in larger data sets.

## 5. Acknowledgements

We would like to thank our patients for their participation in this study and our funding sources: American Brain Tumor Association DG14004, R01CA249882, R01CA218144, R01CA218144-02S1, and R21CA23189201.

---

**Supplemental Figure 1:**
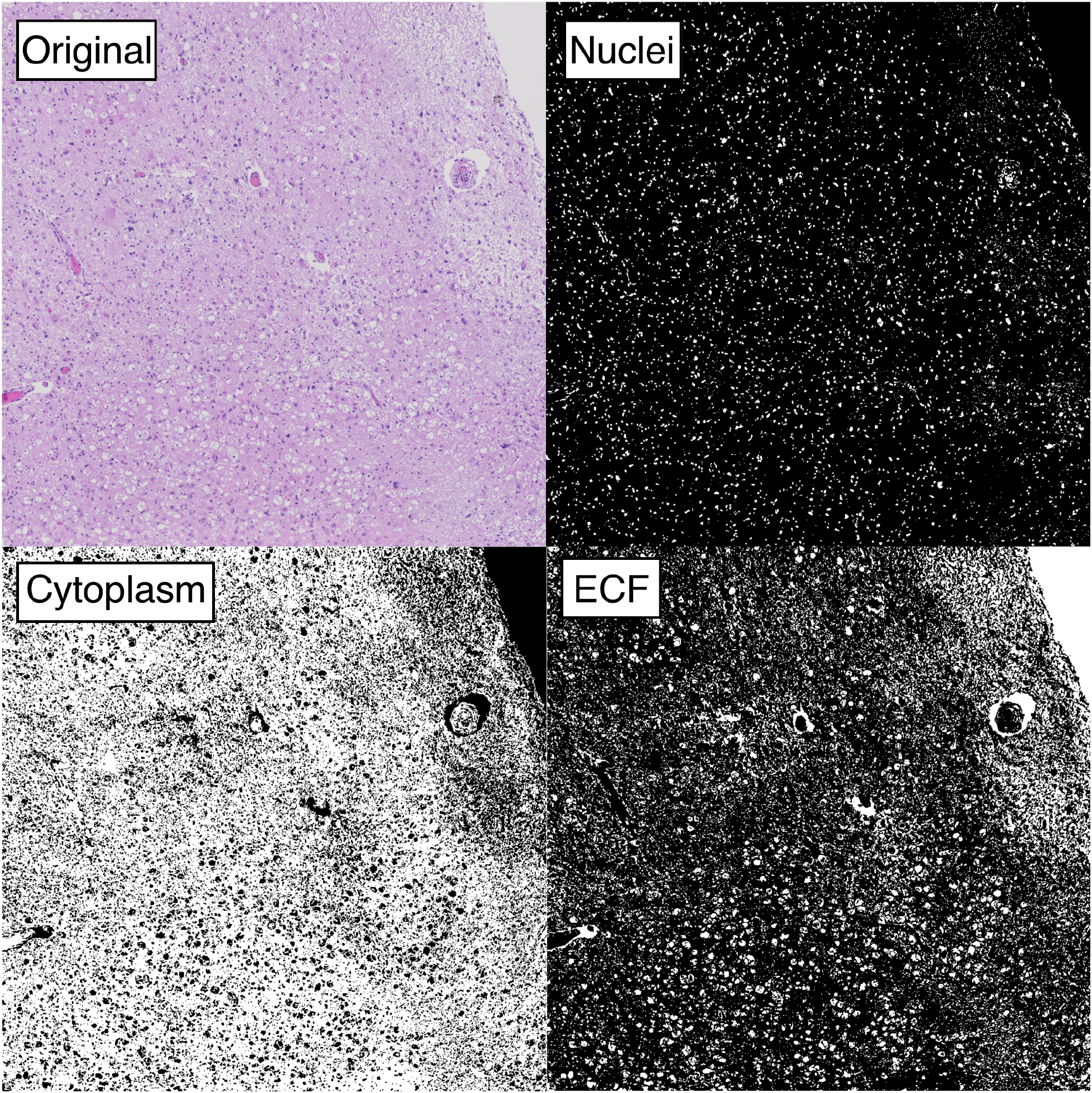
Example tissue segmentations for a representative area of digitized tissue. Segmentations include nuclei (used for computing cell count), cytoplasm, and extracellular fluid (ECF)

**Supplemental Figure 2:**
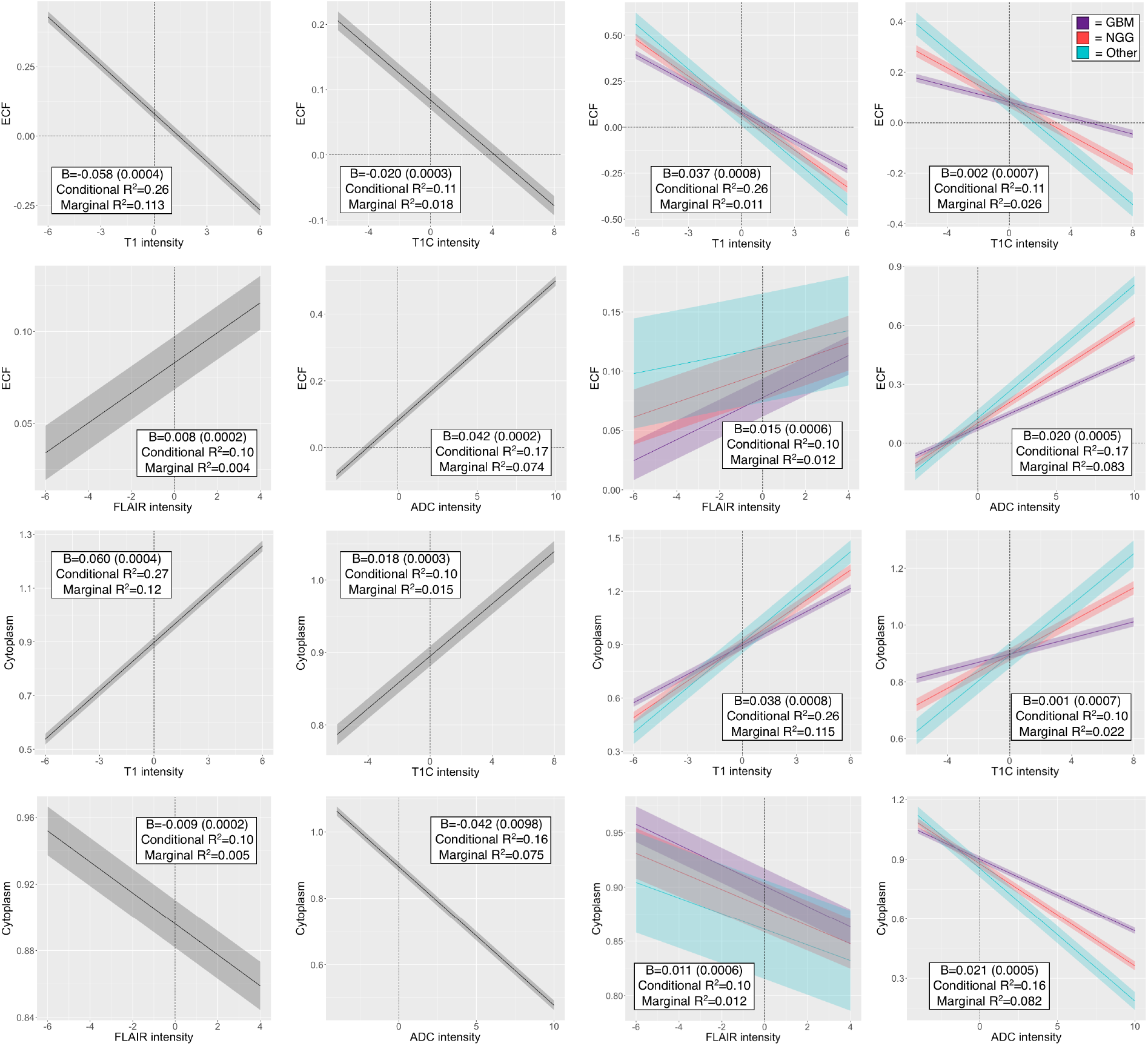
Single image results depicting the relationship between image intensity and ECF/cytoplasm proportion for each contrast. B values for the left-hand plots indicate the change in ECF/cytoplasm proportion per standard deviation increase in image intensity. B values for the right-hand plots indicate the difference in slope between GBM, NGG, and Other patients.

